# Polyamines control inorganic polyphosphate levels during bacterial nitrogen starvation

**DOI:** 10.64898/2026.09.16.752086

**Authors:** Jian Guan, Pavithra Mahadevan, Ursula Jakob

## Abstract

Inorganic polyphosphate (polyP) is a ubiquitous, multifunctional biomolecule that supports stress survival in bacteria. During nitrogen (N) starvation, *Escherichia coli* rapidly accumulates polyP, which drives the formation of RNA–protein granules that sustain long-term survival. What triggers this upregulation has remained unknown. Here we show that polyamines, a group of highly conserved, amino acid-derived polycations, regulate polyP accumulation during N starvation. We found that deleting all nine polyamine biosynthesis genes elevates polyP during exponential growth and by more than 5-fold under N starvation, with shorter chains preferentially accumulating. To dissect which polyamine matters, we further deleted the three catabolic enzymes that consume polyamines as a nitrogen source, which enabled complementation assays. Supplying any one of the three physiological polyamines, putrescine, spermidine, or cadaverine, restored wild-type polyP levels during N starvation. We found that the absence of polyamines did not affect the steady-state levels of either the polyP-synthesizing kinase PPK or the exopolyphosphatase PPX, pointing instead to post-translational control of enzyme activity or to as-yet-unidentified polyamine-dependent regulators. These findings establish a direct metabolic link between two universal, functionally intertwined biomolecules and suggest that the polyamine decline upon N starvation contributes to the observed polyP accumulation.

---

Inorganic polyphosphate (polyP) and polyamines are both highly charged biomolecules present in most cellular organisms (1, 2). PolyP is a linear, nonbranching polymer of covalently linked orthophosphate units, while polyamines are small, linear aliphatic molecules containing two or more amines (1, 2). Despite their chemical distinctions, polyP and polyamines share striking functional overlaps in translation (3-5), bacterial growth (6, 7), swarming motility (8, 9), biofilm formation (10-13) and oxidative stress protection (14, 15). In *Escherichia coli*, polyP is synthesized by the polyP kinase PPK (16, 17) and degraded by the exopolyphosphatase PPX (18) (Fig. 1A). Previous studies showed that polyP is barely detectable under exponential growth conditions but strongly accumulates upon stress conditions, such as nitrogen (N) starvation, osmotic stress and, to a lesser extent, heat shock (19-21). Polyamines, particularly putrescine and spermidine are constitutively present at millimolar levels whereas cadaverine is only detected in stationary phase growth, during low pH, or in the absence of other polyamines (22-24). The biosynthesis of polyamines in *E. coli* involves nine enzymes (Fig. 1A), whose combined deletion causes a slow growth phenotype but no lethality under aerobic conditions (25).

**Figure 1.**
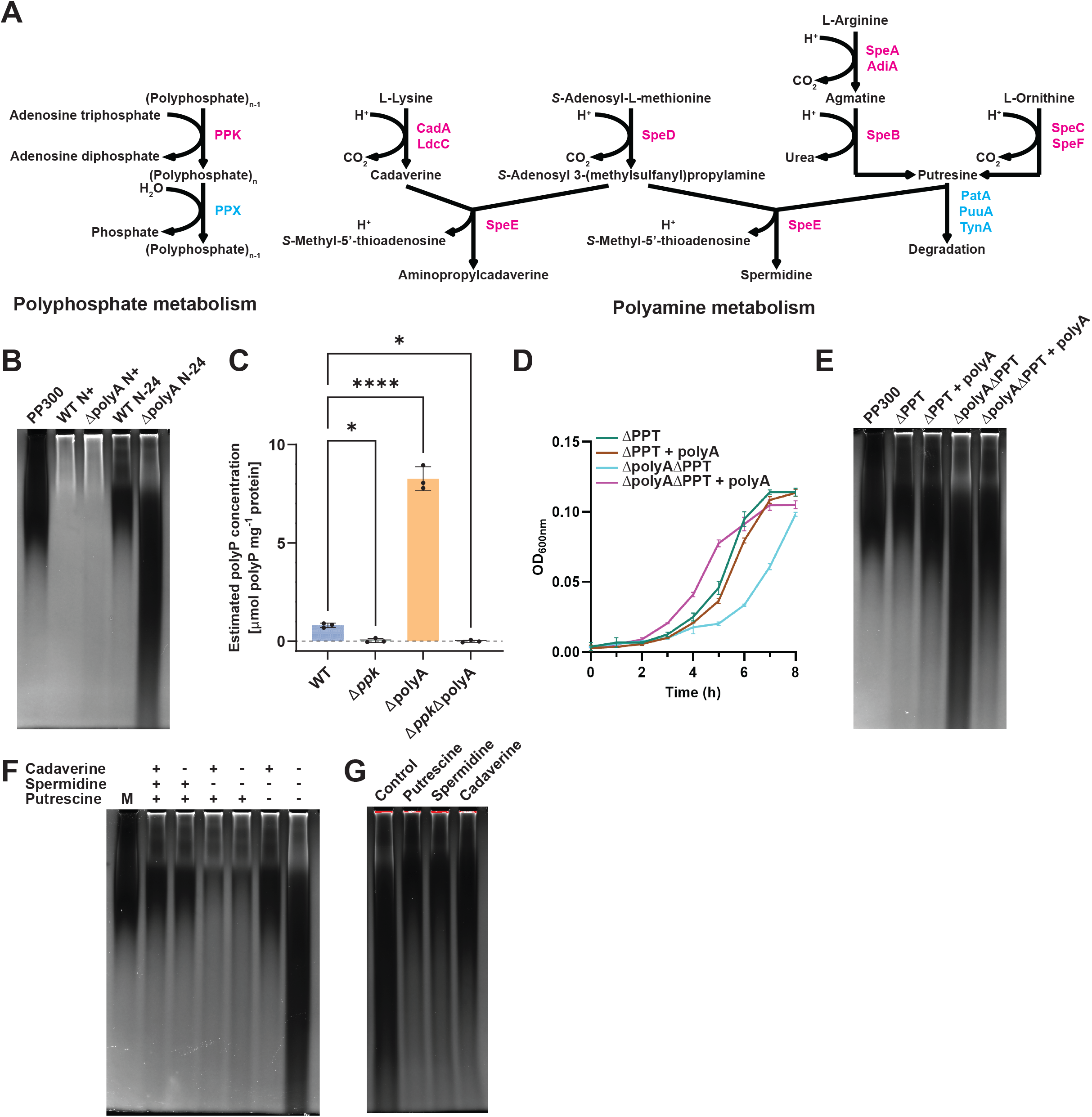
Loss of polyamine enhances polyP accumulation in N-starved *E. coli*. (**A**) Diagram showing major metabolic pathways of polyP and polyamines. Anabolic and catabolic enzymes are shown in magenta and cyan, respectively. Polyamine catabolism is shown only for putrescine for simplicity. (**B**) PolyP extracted from the indicated strains and conditions was visualized on TBE urea gels by DAPI staining and photobleaching. The following strains were used; WT: MG1655; ΔpolyA: Δ*speABBBCBDBEBFB*Δ*adiAB*Δ*cadA*/Δ*ldcC*. PolyP appears as dark patches. Purified polyP 300-mer (chain length of 300 Pi-units) was used as reference (lane 1). (**C**) Quantification of normalized polyP abundance in indicated strains at N-24. One-way ANOVA, *n* = 3, SD, * *p* < 0.05, **** *p* < 0.0001. (**D**) Growth curves of the indicated strains in the presence or absence of polyamine (500 μM putrescine dihydrochloride, 100 μM spermidine trihydrochloride, 500 μM cadaverine dihydrochloride) in the media. (**E)** PolyP extracted from the indicated strains in (D) was visualized on TBE gels by DAPI staining and photobleaching. Purified polyP-300 was used as reference (lane 1). SD, *n* = 3. (**F**) PolyP extracted from strains able (+) or unable (-) to synthesize the indicated polyamines at N-24 was visualized on TBE gel by DAPI staining and photobleaching. The following strains were used: Lane 2: wild-type MG1655; Lane 3: Δ*cadA*/Δ*ldcC* (deficient in cadaverine synthesis); Lane 4: Δ*speD*/Δ*speE* (deficient in spermidine synthesis); Lane 5: Δ*speDEB*Δ*cadA*/Δ*ldcC* (deficient in cadaverine and spermidine synthesis); Lane 6: Δ*speABBBCBFB*Δ*adiA* (deficient in putrescine, which serves as precursor of spermidine); Lane 7: Δ *speABBBCBDBEBFB* Δ *adiAB* Δ *cadA*/Δ *ldcC (*ΔpolyA). Purified polyP-300 was used as reference (lane 1). (**G**) PolyP extracted from ΔpolyA Δppt mutant at N-24 grown in the absence or presence of 2 mM of the indicated polyamines. PolyP was visualized on TBE gel by DAPI staining and photobleaching.

Accumulating evidence suggests that polyP and polyamines may reciprocally affect each other’s cellular abundance. An earlier study reported that partially disrupting polyamine biosynthesis in *E. coli* reduces polyP accumulation during acute amino acid depletion (26). More recently, an engineered *Citrobacter freundii* strain overproducing polyP was shown to accumulate more spermidine and spermine, likely due to elevated expression of several polyamine-producing enzymes (27). These findings suggest a potential crosstalk between biosynthetic pathways of the two oppositely charged biomolecules.

To investigate whether the polyP accumulation that we observed within 24h of N starvation in *E. coli* (21) was at all connected to bacteria depleting their polyamine stores, we constructed an *E. coli* MG1655 strain deleted for all nine genes that participate in polyamine biosynthesis (i.e., ΔpolyA-strain) (25). Whole genome sequencing of this ΔpolyA strain revealed chromosomal rearrangements and the loss of a 20 kb fragment during its construction (Fig. S1) that were similar to previous reports (28). When we grew wild-type and the ΔpolyA using our previously established N starvation regimen (21) and extracted the polyP, we found that the ΔpolyA strain not only accumulated significantly higher levels of polyP but with a chain length distribution that spanned a much wider range compared to the WT strain (Fig. 1B). Of note, we also noticed that the ΔpolyA strain accumulated detectable amounts of short-chain polyP under N-replete conditions (N+), suggesting that the lack of polyamine promotes polyP synthesis and/or prevents polyP hydrolysis (Fig. 1B). Biochemical quantification confirmed our results on TBE gels and revealed a more than 5-fold increase in polyP levels in N-starved *E. coli* lacking the polyamine biosynthetic machinery compared to WT *E. coli* (Fig. 1C). These results provide evidence for a hitherto unknown connection between polyamine and polyP levels in *E. coli*.

To test the possibility that the chromosomal rearrangements that arose during the stepwise construction of the 9-gene deletion strain ΔpolyA contribute to the observed effects, we considered complementation assays using supplementation with putrescine, spermidine and/or cadaverine in the media. We reasoned that if the effects were indeed mediated by the lack of these polyamines, we should, at a minimum, restore WT-levels of polyP in the polyA strain.

However, since N-starved bacteria use exogenously administered polyamine as an alternative N source (29), we needed to first modify our strains in a way that they were no longer able to metabolize polyamines. We therefore deleted the three genes, i.e., *patA, puuA, tynA* that are known to catalyze polyamine catabolism in *E*.*coli* (29, 30) in the background of an otherwise wild-type strain, generating the ΔPPT strain, or in the background of the ΔpolyA strain, generating the 12-gene deletion strain ΔpolyA/ΔPPT. Importantly, we found that the addition of all three polyamines (i.e. polyA) to the growth medium effectively rescued the growth defect of the ΔpolyA/ΔPPT without benefiting the ΔPPT strain (Fig. 1D). Even more importantly, however, addition of the polyamines eliminated the polyP over-accumulation under N starvation, effectively bringing the polyP levels back to WT levels (Figs. 1E). Together, these data suggested that the presence of polyamine in *E. coli* either directly or indirectly contributes to a reduction in polyP levels and that their depletion enhances polyP accumulation upon N-starvation.

To understand whether the different polyamines show distinct impacts on cellular polyP level, we next constructed *E. coli* strains deficient in either the synthesis of the individual polyamines or possible combinations thereof. Neither deleting the two enzymes catalyzing the synthesis of cadaverine (i.e., CadA, LdcC) nor the five enzymes synthesizing putrescine (i.e., SpeA,B,C,F/AdiA) caused any detectable increase in polyP beyond wild-type levels (Fig. 1F). While the first result was consistent with the low abundance of cadaverine under normal growth conditions, the second result could be explained by the compensatory upregulation of cadaverine when putrescine is missing (25). Of note, however, deleting the genes *speD* and *speE*, which encode for enzymes that convert putrescine into spermidine, either alone or in combination with mutants of the cadaverine synthesis pathway reduced polyP to levels that were substantially lower than in N starved wild-type *E. coli* (Fig. 1F). Supplementation of the ΔpolyA ΔPPT strain with either one of the three polyamines restored wild-type like polyP levels in N-starved *E. coli* to a similar extent, providing evidence that the individual polyamines indistinguishably suppress the over-accumulation of polyP (Fig. 1G). These results suggest a specific role for SpeD/E in polyP regulation that is independent of the product spermidine and support the conclusion that the absence of all three polyamines is necessary to cause the massive over-accumulation of polyP upon N-starvation.

To test whether the over-accumulation of polyP in the ΔpolyA strain was due to changes in the abundance of the PPK or PPX enzyme, we made cell lysates of N-starved WT and ΔpolyA strains and conducted Westernblot analysis. As shown in Figs. 2A-B, we did not observe any significant difference in the steady-state levels of either enzyme. These findings were consistent with earlier observations that PPK expression does not correlate with polyP production (31, 32). Moreover, and based on our previous results that deletion of *ppx* only slightly increases polyP levels under N starvation (21), we also excluded the possibility that inactivation of the exopolyphosphatase in the absence of polyamine might be responsible for the observed polyP increase. To test whether polyamines inhibit the enzymatic activity of PPK, which would explain the increase on polyP levels in their absence, we conducted *in vitro* polyP synthesis assays with purified *E. coli* PPK in the absence and presence of putrescine. Consistent with the ability of polyamines to act as chemical chaperones (33), we found that addition of putrescine slightly stimulates the polyP-synthesizing activity of PPK (Fig. 2C), which, if relevant *in vivo*, would cause a decrease and not the observed increase in polyP levels when polyamines are absent. These results suggest that polyamine’s effect on N starvation-induced polyP accumulation might involve additional proteins or components that affect the polyP biosynthetic machinery in *E. coli*.

**Figure 2.**
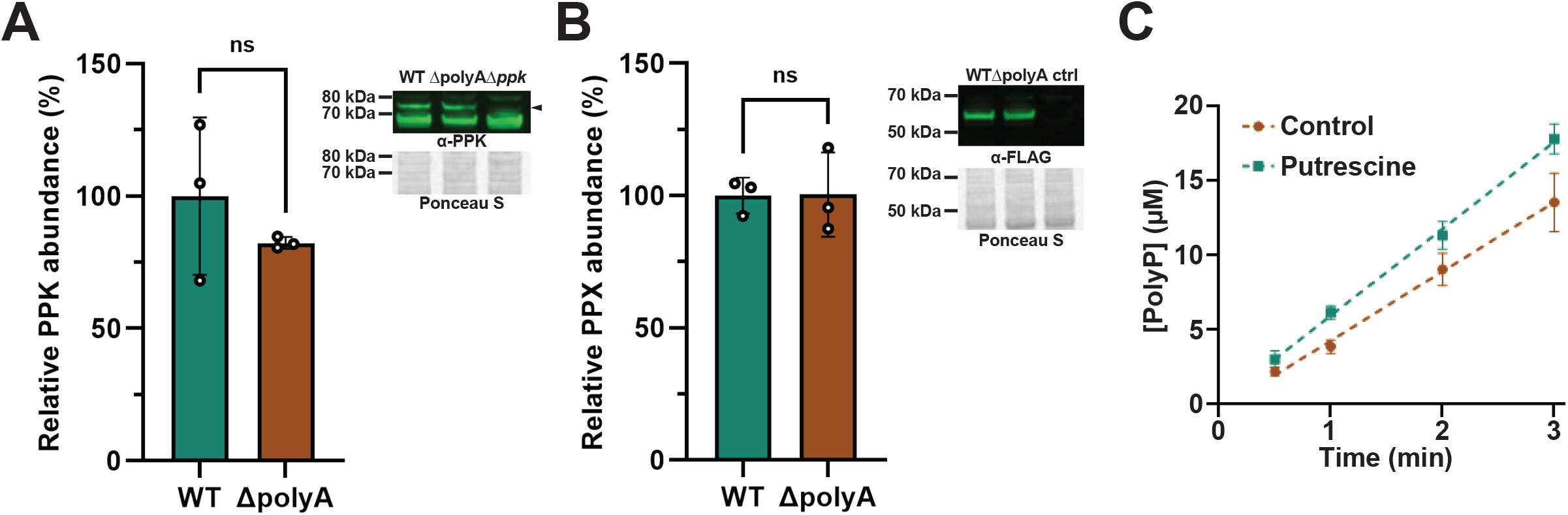
Polyamine deficiency does not alter PPK/PPX abundance. (**A**) Protein abundance of PPK in WT and ΔpolyA strains at N-24 as quantified by Westernblot analysis using antibodies against endogenous PPK. A Δ*ppk* strain was used as negative control. The PPK band is shown by the arrowhead. Total protein was used as loading control. *t*-test, *n* = 3, SD. (**B**) Protein abundance of endogenously expressed PPX-3×FLAG in WT and ΔpolyA strains at N-24 as quantified by Westernblot analysis using antibodies against the FLAG-tag. An untagged WT strain was used as negative control. Total protein was used as loading control. *t*-test, *n* = 3, SD. (**C**) *In vitro* polyP synthesizing activity of purified *E. coli* PPK (5 nM) in the absence or presence of 10 mM putrescine. SD, *n* = 3.

In summary, we show a hitherto unknown interaction between two ancient and highly conserved biosynthetic pathways. *E. coli* has long served as the canonical model for polyP metabolism, yet a central paradox has remained unexplained: exponentially growing cells contain little polyP even when the hydrolyzing enzyme PPX is deleted (31, 32), PPK expression levels do not correlate with polyP levels and yet, specific stress conditions trigger robust polyP accumulation. Polyamines might offer a solution to this conundrum. Polyamine levels fall as growth rate and nutrient availability decline (34) and bacteria mobilize polyamines as an alternative nitrogen source during N limitation (35, 36). Under many of the same circumstances, polyP levels have been shown to increase (19). We now propose that under N-replete non-stress conditions, bacteria contain millimolar concentrations of polyamines, such as putrescine that actively restrain polyP levels by suppressing its synthesis, stimulating its hydrolysis, or both. This model would also explain why ΔpolyA cells accumulate polyP even during exponential growth.

Mechanistically, we consider the possibility that direct polyamine–polyP interactions may alter how PPK and PPX engage with polyP, tipping the balance away from synthesis and toward degradation. Alternatively, *E. coli* may harbor as-yet-unidentified polyphosphatases or kinases whose activity is either directly or indirectly controlled by polyamines. Either way, the logic stays the same: as polyamines are depleted during N starvation, they can no longer suppress polyP, and polyP accumulates. The disproportionally higher polyP accumulation of the ΔpolyA strain under N-deplete compared to N-replete conditions furthermore suggests that additional polyamine-dependent systems regulate polyP synthesis specifically during N starvation. Future work will reveal how polyamines govern polyP homeostasis under non-stress and stress conditions, and how far this regulation extends on an evolutionary scale.

## Material and Method

### Media and culture conditions

All bacterial growth assays, unless stated otherwise, were performed in culture tubes at 37°C with 200 rpm agitation. To induce nitrogen (N) starvation, bacteria were grown overnight in N+ Gutnick medium (33.8 mM KH_2_PO_4_, 77.5 mM K_2_HPO_4_, 5.74 mM K_2_SO_4_, 0.41 mM MgSO_4_, 0.4% w/v glucose, 10 mM NH_4_Cl, 10 μM ferric citrate and micronutrients (3 nM (NH_4_)_6_Mo_7_O_24_, 400 nM H_3_BO_3_, 30 nM CoCl_2_, 10 nM CuSO_4_, 80 nM MnCl_2_, 10 nM ZnSO_4_)(34, 35). Overnight cultures were diluted with and grown in N-Gutnick medium (same as N+ Gutnick medium but only 3 mM NH_4_Cl). Bacteria enter N starvation within 6h of growth at 37°C (21, 36). Cells were analyzed either during mid-log phase (N replete conditions, N+) or 24h after entering N starvation (N24). Luria-Bertani (LB) broth and Super Optimal Broth with Catabolite repression (SOC) medium were used for all cloning experiments.

### Bacterial strain construction

Bacterial strains and plasmids used in this study are listed in Table S1. PCR amplifications were performed with Platinum Superfi II Polymerase (Thermo Fisher Scientific). Primers used in this study are listed in Table S2. Deletion of *speAB* and *speDE* and chromosomal FLAG-tag addition were performed using the Datsenko-Wanner method employing λ-Red recombination (37). Briefly, linear DNA flanked by 50 bp homology arms were electroporated into competent *E. coli* str. K-12 substr. MG1655 carrying pKD46 plasmid. Recombination was selected by kanamycin resistance. Positive colonies were confirmed by PCR screening and sequencing (Plasmidsaurus). Other gene deletions were performed with P1 transduction using donor strains from the Keio collection (38), and positive colonies were validated by PCR screening. To cure selection markers, pCP20 plasmids were electroporated into competent bacteria, and expression was induced at 43°C. Kanamycin-sensitive cells were selected for further experiments.

### PolyP extraction, quantification and visualization

Cells were grown under N+ conditions or N starved for 24h. Then, 2 ODmL bacterial culture were collected and resuspended in 400 μl chilled AE buffer (50 mM sodium acetate pH 5.3, 10 mM EDTA, pH 8.0). Samples were sonicated on ice with 70% power for six seven-second cycles. 5 μL lysate was collected and mixed with 15 μL GITC buffer (4 M guanidium isothiocyanate, 50 mM Tris-HCl pH 7.0) for protein quantification using a BCA assay (Thermo Fisher). 300 μL phenol and 40 μL 10% w/v SDS were added and the vortexed mixture was incubated at 65°C for 5 min, and then on ice for 2 min. After addition of 300 μL chloroform, the mixture was vortexed and centrifuged at 4°C, 13,000 g for 2 min. The upper aqueous phase was removed and mixed with 350 μL chloroform, vortexed and centrifuged again. After collecting the aqueous phase and measuring the sample volume, 3-times the volume of 100% ethanol, 10% volume of 3 M sodium acetate, pH 5.3 and 2 μL of a 5 mg/mL glycogen solution (Thermo Fisher) were added. The vortexed mixture was incubated on ice for 1 hour. Nucleic acids, polyP and glycogen were pelleted at 4°C, 17,000 g for 20 minutes, washed once with chilled 70% ethanol and air dried. Dried pellets were dissolved in 100 μL B buffer (50 mM Tris-HCl pH 8.0, 20 mM NaCl, 2 mM MgCl_2_). 0.1 μL Benzonase (Sigma-Aldrich) was added to each sample and nucleic acids were digested at 37°C for 1.5 hours. The enzyme was removed by chloroform extraction. PolyP abundance was quantified as previously reported (39). Briefly, 1 μL purified yeast PPX (40) was added to 30 μL Benzonase-digested sample and incubated for 2 hours at 37°C. To measure the release of free phosphates, assay reagents were freshly prepared by mixing 91.2 (v/v) parts of reagent A (2.4 mM ammonium heptamolybdate; 600 mM sulfuric acid, 0.6 mM potassium tartrate) and 8.8 parts (v/v) of reagent B (88 mM Ascorbic acid). 75 μL of this reaction mixture was added to 25 μL of sample in 96-well half area plates (Corning) followed by a 5-minute incubation at room temperature. Absorbance at 882 nm was read in a plate reader (Tecan Infinite 200 Pro) and concentrations were calculated based on a standard curve generated with known concentrations of potassium phosphate solution. To visualize polyP chain length distribution and semi-quantitatively assess polyP abundance, 10 μL Benzonase-digested polyP sample was mixed with 2 μL DNA gel loading buffer. 10 μL of the mix was loaded onto a 8% Novex TBE gels (Thermo Fisher). 10 μL 500 μM polyP 300-mer was loaded as marker. Gel electrophoresis was performed on ice in TBE buffer (89 mM Tris base, 89 mM boric acid, 2 mM EDTA) at 100 V for 80 minutes. The gel was stained with 2 μg/mL DAPI in 25% methanol, 5% glycerol and 50 mM Tris base with agitation for 30 minutes and then washed for 30 minutes in the same buffer without DAPI. The DAPI-stained gel was imaged on Chemidoc and polyP was visualized after 8 minutes UV photobleaching.

### Growth curve measurement

Bacterial growth in N-Gutnick medium was measured by collecting 100 μL culture every hour into transparent 96 well plates (Corning). OD at 600 nm was obtained on a TECAN M1000 plate reader. Fresh N-Gutnick medium was used for background subtraction.

### Immunoblotting

To quantify protein abundance, 0.3 ODmL cultures were collected, washed and resuspended in 30 μL SDS-polyacrylamide gel loading buffer (6.5 mM Tris-HCl pH 7.0, 10% glycerol, 2% SDS, 0.05% bromophenol blue, 2.5% β-mercaptoethanol). Samples were incubated at 95°C for 10 minutes and loaded onto a 4-12% NuPAGE Bis-Tris gel (Invitrogen). After running at 175V for 45 minutes, the gel was transferred to a polyvinylidene difluoride membrane (Bio-Rad) and blocked with EveryBlot blocking buffer (Bio-Rad) for 5 minutes with agitation. The membrane was then incubated at RT in blocking buffer containing 1:1000 diluted mouse anti-FLAG M2 monoclonal (Sigma) or 1:500 rabbit anti-PPK polyclonal antibody for one hour. After five washes with TBST, the membrane was incubated at RT in blocking buffer containing 1:10000 diluted goat anti-mouse 800CW (LI-COR Biosciences) or goat anti-rat HRP secondary antibody for one hour. Following three washes in TBST, the membrane was imaged with LI-COR Odyssey CLx imager or Chemidoc using SuperSignal West Pico PLUS (Thermo Fisher). Gel quantification was performed with ImageJ, and normalization was performed against total protein by Ponceau S staining of the same blot prior to blocking. Three biological replicates were included in all experiments.

### *In vitro* assay of PPK activity

PPK activity was determined as previously described (32). Briefly, 125 μL reaction mixtures containing 5 nM C-tagged *E. coli* PPK (41), 50 mM HEPES-KOH (pH 7.5), 50 mM (NH_4_)_2_SO_4_, 5 mM MgCl_2_, 20 mM creatine phosphate and 60 μg/mL creatine kinase with and without 10 mM putrescine dihydrochloride were prepared. Mixtures were prewarmed to 37°C, and reactions were initiated by adding MgATP to a final concentration of 6 mM. To detect polyP, 20 μL aliquots were taken from the reaction mixtures at 0.5, 1, 2 and 3 min and the reaction was quenched by diluting the sample into 80 μL of 62.5 mM EDTA-50 μM DAPI in black 96-well plates. PolyP-DAPI fluorescence (excitation 415 nm, emission 600 nm) was measured with TECAN M1000 microplate reader. PolyP concentration (calculated in terms of individual phosphate monomers) was determined by comparison to a 0-150 μM standard curve of polyP 300-mer, and rates of polyP synthesis were calculated by linear regression.

### Statistical analysis

All statistical analyses were performed using Graphpad Prism and detailed in corresponding figure legends.

## Acknowledgements

We thank M. Gray for providing us with purified PPK enzyme and T. Shiba for providing polyP-300. This work was supported by NIH grant GM122506 to U.J.

