## Supplemental Information for "Polyamines control inorganic polyphosphate levels during bacterial nitrogen starvation"

**SUPPLEMENTAL MATERIAL**

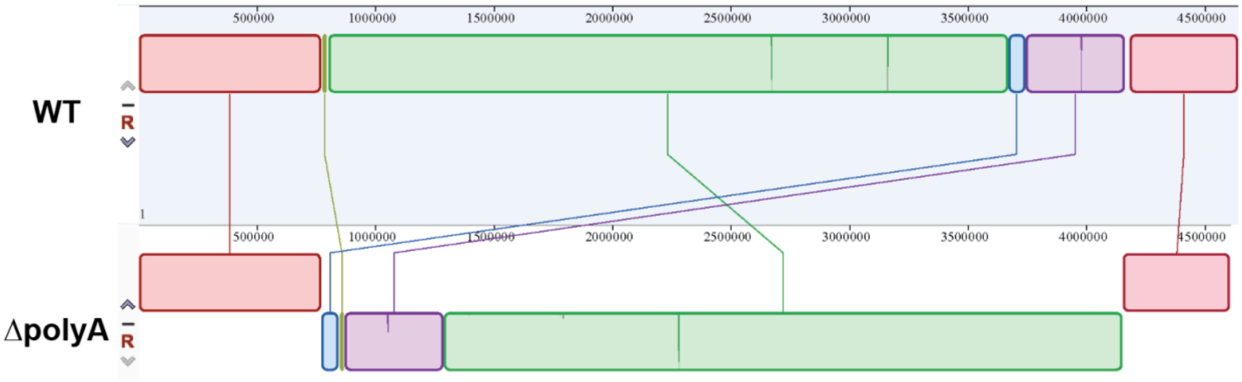


**Figure S1. Genomes of indicated *E. coli* MG1655 strains.** Chromosomal rearrangements are visualized with Geneious Prime. A 20 kb fragment between *adiA* and *cadA* is lost during Flp-mediated recombination. No essential genes are present in this region.

**Table S1 Bacterial strains and plasmids used in this study**

| **Bacterial strain** | | **Marker** | **Source** |
|---|---|---|---|
| *E. coli* str. K-12 substr. MG1655 (wildtype) | | | Lab collection |
| MG1655 Δ*adiA* Δ*cadA* Δ*ldcC* Δ*speAB* Δ*speC* Δ*speDE* Δ*speF* (ΔpolyA) | | | This study |
| MG1655 *speDE::Kan* | | Kan | This study |
| MG1655 Δ*cadA ldcC::Kan* | | Kan | This study |
| MG1655 Δ*cadA* Δ*speDE ldcC::Kan* | | Kan | This study |
| MG1655 Δ*speAB* Δ*speC* Δ*speF adiA::Kan* | | Kan | This study |
| MG1655 Δ*ppk* | | | (1) |
| MG1655 Δ*adiA* Δ*cadA* Δ*ldcC* Δ*speAB* Δ*speC* Δ*speDE* Δ*speF ppk::Kan* | | Kan | This study |
| MG1655 Δ*patA* Δ*puuA tynA::Kan* (ΔPPT) | | Kan | This study |
| MG1655 Δ*adiA* Δ*cadA* Δ*ldcC* Δ*speAB* Δ*speC* Δ*speDE* Δ*speF* Δ*patA* Δ*puuA tynA::Kan* | | Kan | This study |
| MG1655 *ppx::ppx-3×FLAG* | | Kan | This study |
| MG1655 Δ*adiA* Δ*cadA* Δ*ldcC* Δ*speAB* Δ*speC* Δ*speDE* Δ*speF ppx::ppx-3×FLAG* | | Kan | This study |
| **Plasmid** | **Description** | **Marker** | **Source** |
| pSUB11 | FLAG tag donor | Kan | (2) |
| pKD46 | λ-red recombinase expression plasmid | Amp | (3) |
| pCP20 | Flp recombinase expression plasmid | Amp | (3) |

**Table S2 Primers used in this study**

| Primer | Sequence |
|---|---|
| adiA-F | GAAGATACTTGCCCGCAACGAAGATTC |
| adiA-R | GAATCCAGGCGAATTCATCGACAAGCTC |
| ldcC-F | GTTTGAGCAGGCTATGATTAAGGAAG |
| ldcC-R | CATCCATCGTCGAGTGGGTGTTGATGAAG |
| cadA-F | CAGAGCCACTCAATGGATAACCACAC |
| cadA-R | CATCGAGAGTGGAATACGTATTAGCGAAC |
| speC-F | GAGCTGGTGACCAGTTTGACCCATATCTCATG |
| speC-R | CAGGTAATTCAACAGCATGTTCGGAATAC |
| speF-F | CACGATCGATTTCTCATTCGAGAAATTGAG |
| speF-R | CTGTTACCTTGATTGACATAATCGACCAGTG |
| speAB-in-F | CACTTAATAAAATAATTTGAGGTTCGCTCGATTGTGTAGGCTGGAGCTGCTTC |
| speAB-in-R | TACCCGTGCGCATCGCATCTGGTGCGAATATCCTCCTTAGTTCCTATTC |
| speAB-out-F | GCGATAGTCGTTAACTGTTTTACACTTAATAAAATAATTTGAGGTTCGCTC |
| speAB-out-R | CGCATCCGACATTAATGGCACGTTTTACCCGTGCGCATCGCATCTG |
| speAB-F | CACTGCTGGAAAATCCATGTGCTTATG |
| speAB-R | CGAACGTAGGTCAGATAAGGCGTTC |
| speDE-in-F | GTGTTAACAAAGGAGGTATCAACCCCGATTGTGTAGGCTGGAGCTGCTTC |

| | |
|---|---|
| speDE-in-R | GAGCCTGGGAGCTCCGCCAGAGCCGGAATATCCTCCTTAGTTCCTATTCC |
| speDE-out-F | ATTATGTTGCGCCCTTTTTTTACGGGTGTTAACAAAGGAGGTATCAAC |
| speDE-out-R | GCAGCAGAAGTAAATAAATCTGGCGGAGCCTGGGAGCTCCGCCAGAG |
| speDE-F | CTACGTCAAATAATCCCTGATAC |
| speDE-R | GATGAACTACTACGACGAAGAAAG |
| patA-F | CATTTGCGCAGCAATCATCAAATC |
| patA-R | GTGGTGGTCTTATGCAATCTG |
| puuA-F | GTGGACTAAATTATCGCCATTACTG |
| puuA-R | TCATCGGCATCATCTCATTTCCTC |
| tynA-F | GTGACGTTGTCACATTATGCATG |
| tynA-R | CATACCTTAATTGACGCTAACTTAC |
| ppx-F-in-F | AGAAAGTACACCAGAAATCGCCGCTGACTACAAAGACCATGACG |
| ppx-F-in-R | ACATTTCTCGTCGGCCCGCAAAGTACCTCCTTAGTTCCTATTCC |
| ppx-F-out-F | CTGGCTGGCGGTTGAAAATTGAAGAAGAAAGTACACCAGAAATCG |
| ppx-F-out-R | GAAAGTGCCTGAATAATGCGGGCCGACATTTCTCGTCGGCCCGCAAAG |
| ppx-F-seq-F | CAGTTCCTGCCACTGATACAGCTATTG |
| ppx-F-seq-R | GCCCAGTAGCTGACATTCATC |


1. Beaufay F, Amemiya HM, Guan J, Basalla J, Meinen B, Chen ZY, Mitral R, Bardwell JCA, Biteen JS, Vecchiarelli AG, Freddolino PL, Jakob U. 2021. Polyphosphate drives bacterial heterochromatin formation. Science Advances 7.
2. Uzzau S, Figueroa-Bossi N, Rubino S, Bossi L. 2001. Epitope tagging of chromosomal genes in Salmonella. Proceedings of the National Academy of Sciences of the United States of America 98:15264-15269.
3. Datsenko KA, Wanner BL. 2000. One-step inactivation of chromosomal genes in Escherichia coli K-12 using PCR products. Proceedings of the National Academy of Sciences of the United States of America 97:6640-6645.
